# [mixglm]: A tool for modelling ecosystem resilience

**DOI:** 10.1101/2024.03.23.586419

**Authors:** Adam Klimeš, Joseph Daniel Chipperfield, Joachim Paul Töpper, Marc Macias-Fauria, Marcus Spiegel, Vigdis Vandvik, Liv Guri Velle, Alistair William Robin Seddon

## Abstract

**Aims:** A number of modelling frameworks exist to aid in the identification and exploration of stable states and the assessment of resilience from ecological datasets. However, because such models are complex to implement there is a substantial barrier for the application in ecological research. Here we develop a flexible model of ecological resilience based on Bayesian approximation of the “stability landscape”. We illustrate its usage on a tropical area where variation in tree cover has been previously interpreted as alternative stable states.

**Methods:** The stability landscape, from which stable states and resilience parameters are computed, is modelled using a mixture of multiple distributions, each representing a regression between the system state variable and the environmental covariates. Our “mixglm” model allows the mean, precision, and probability parameters of these distributions in the landscape to be dependent on multiple external covariates. “Mixglm” is implemented as a function in R package with the same name, internally using Bayesian inference via NIMBLE. We also conducted a power analysis to provide guidance regarding required sample size.

**Results:** We illustrate the use of the “mixglm” on a published case of tree cover in South America which reports a stability landscape with three distinct stable states. Using “mixglm”, we were able to replicate the identification of these states. Moreover, we quantified uncertainty of our estimates, and computed resilience of South America’s forests.

**Conclusions:** “Mixglm” can be readily used for description of stability landscapes and identification of stable states in most spatial datasets of system state variables, and it is accompanied by tools for calculation of resilience metrics. It can also be further expanded using regression framework to account for more complex data structures such as spatio-temporal data.

## Introduction

Under certain conditions, ecological systems exist as multiple alternative stable states separated by thresholds or tipping points (Pausas & Bond, 2020; Scheffer et al., 2001). Transitions between these states can result in large differences in ecological structure and functioning. Because transitions between alternative states can occur abruptly and can be to difficult reverse, there is global concern about the status of multiple Earth system components and their potential for abrupt change from the local to the global scale (Armstrong McKay et al., 2022). To avoid or adapt to these changes, information about stable states of systems, their alternative equilibria and the associated ecological thresholds is of primary importance.

According to theory, stable states are maintained by internal negative feedbacks between different components of the system. These feedbacks counterbalance any deviations from a stable system equilibrium following an external disturbance (Holling, 1973) or through internal stochastic forcing (Scheingross et al., 2020). Under certain conditions, disturbance or internal stochastic dynamics can induce a series of positive feedbacks, reducing the ability of the system to return to its original state and resulting in a transition to an alternative stable state (Scheffer et al., 2009, 2012).

The concept of ecological resilience provides an overarching framework for such ideas. Resilience and its associated terms (e.g.: resistance, precariousness; Hodgson et al., 2015) generally describe the ability of a system to resist the change and/or recover from disturbance (Hodgson et al., 2015; Yi & Jackson, 2021). To preserve ecosystem functions, increase of ecosystem resilience was set as one of the targets for global biodiversity management (target 15; Convention on Biological Diversity, 2010). However, although resilience research is prominent in the literature, there are varied definitions and methods used to quantify resilience using ecological datasets. This presents a challenge for system assessment and priority mapping because metrics derived between different studies are often not comparable.

Stable states, although often marked as attractors of a system, are result of interactions between a variety of system components such as plant growth, herbivory consumption or fire. Separated, these components “push” the system to state values which can be (and typically are) distinct from state values of stable states (Fig. 1A). When effects of these components cancel each other out, the system is in equilibrium. Equilibria can either be stable (i.e. representing a stable state) or unstable (i.e. constituting a tipping point; Holling, 1973). Such a system can be visualized using a stability curve with valleys representing stable states and hilltops representing tipping points (Fig. 1B; Hodgson et al., 2015; van Nes et al., 2016). A change in the stability curve along an external (e.g., climatic) gradient then constitutes the stability landscape (Fig. 1C; *sensu* Scheffer et al., 2001). The existence of stable states is often dependent on these external conditions (Scheffer & Carpenter, 2003).

**Figure 1:**
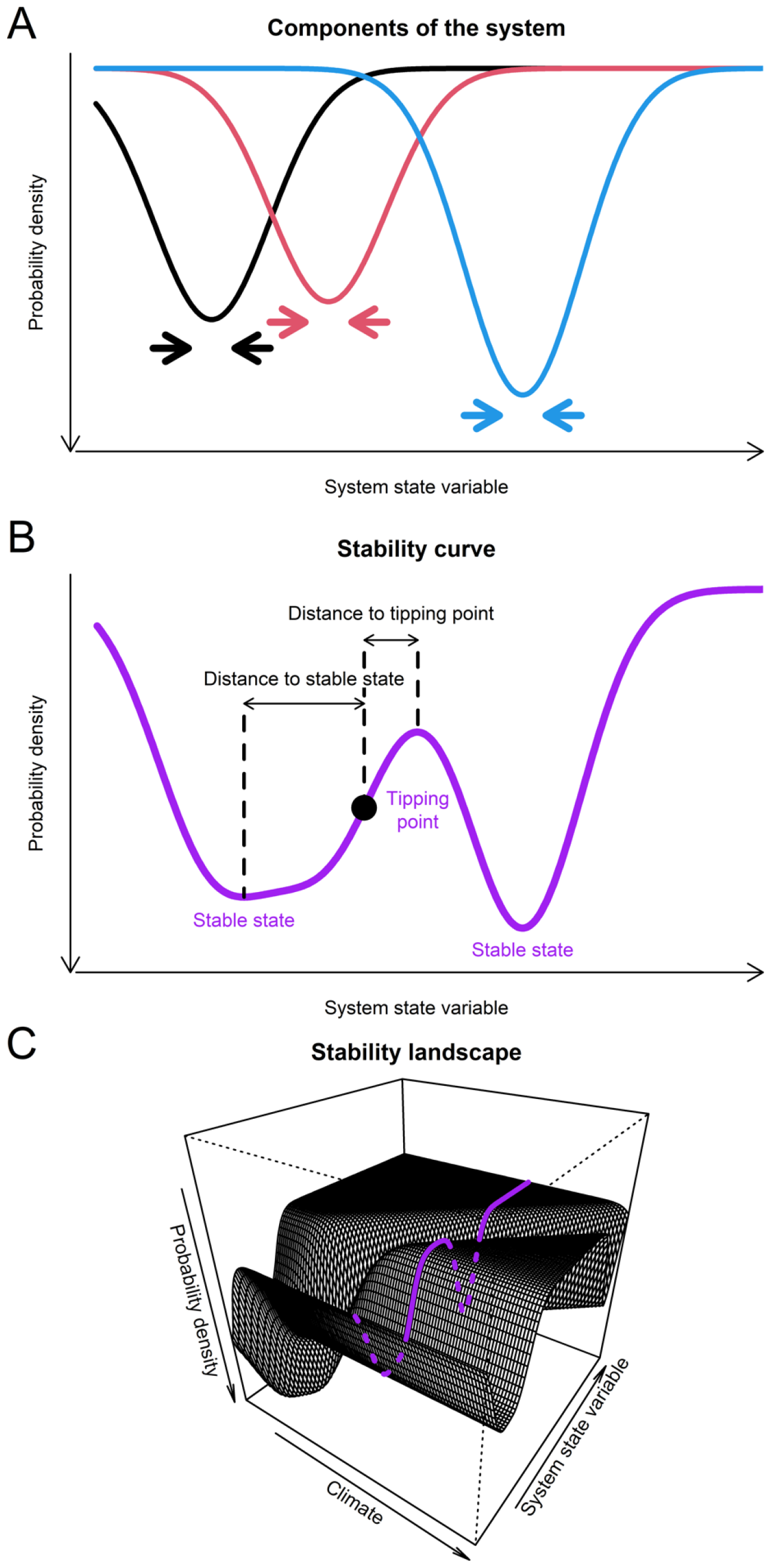
Components of the system, stability curve, and stability landscape. A: Visualization of effects of three components of the system which “push” the system state value in direction marked by arrows. B: Stability curve which results as a combination of effects of the three system components. Although there are three components, the system has only two stable states. Black point denotes an instance of the system. C: Simulated stability landscape. Stability curve can change along climatic gradient resulting in stability landscape. Purple line highlights stability curve from Fig. 1B.

According to this framework, a system with a wide stable state basin is typically capable of withstanding large disturbances without experiencing a transition to an alternative state (i.e. it has large ecological resilience *sensu* Gunderson, 2000). In contrast, systems with narrow stable state basins are prone to changes to alternative states. Finally, steep slopes indicate that a stable state has high resistance to a change and/or a high recovery rate. There is no way to distinguish between resistance and recovery rate of a stable state in this framework, as high values of both result in steep slopes of the stability curve (this contrasts with so called “classical resilience landscape” as described by Hodgson et al., 2015 which does not capture resistance at all).

Whilst the theoretical ideas that underly the presence and location of alternative stable states has long been recognised, practical application to identify alternative states in empirical datasets remains a challenge (Scheffer & Carpenter, 2003). One set of approaches have used time series analyses to identify abrupt changes in a state variable (e.g., Smith et al., 2022), or to identify changes in the statistical properties of a state variable which may provide clues of an imminent transition to an alternative state (e.g., Feng et al., 2021; Forzieri et al., 2022; Rocha, 2022). Serious limitation of these approaches is the length of available time series which for most remote sensed products does not exceed a few decades (e.g., Moesinger et al., 2020). An alternative approach involves using the distribution density of a state variable in relation to external predictor(s) to describe the system states (Hirota et al., 2011; Scheffer et al., 2012). Here, areas in the state variable space along a predictor variable with high probability density are assumed to suggest the presence of a stable state. Such an approach presents a pragmatic method to infer system stable states based on spatial dataset (without timeseries data), from which properties future dynamics and transitions among states can be predicted. However, due to model complexity, this approach is rarely used or is not used to its full potential (e.g., using none or only one external predictor; Hirota et al., 2011).

Here, we develop a model which can parametrize stable states using a mixture of distributions, which represent putative components of the system. By making these distributions dependent on external predictors, we can then parametrize the whole stability landscape. To our knowledge, at present there is no available implementation of this type of model and the absence of a readily available tool is a limiting factor for ecological resilience research. We implement our model as a function in R package “mixglm” (https://github.com/adamklimes/mixglm; R Core Team, 2023), which internally uses Bayesian inference via NIMBLE (de Valpine et al., 2017). We illustrate its use on tree cover in the tropical region of South America, which has previously been suggested to have three distinct stable states – forest, savanna, and treeless state (Hirota et al., 2011) and we run power analyses to provide guidance regarding required sample size.

## Material and Methods

### Model

For a system with one component (and thus one stable state), resilience can be modelled by a regression with the type depending on the distribution of system state values (e.g., gamma regression). Furthermore, the strength of the effect of the component may change along external predictors (such as climate). Therefore, we want not only the mean but also the variance/precision to be dependent on these predictors. Such a model for a normally distributed system state variable (y) is (using matrix notation):

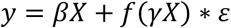

where β and γ are vectors of parameters for mean and variance respectively, X is the matrix of predictors, f is a link-function (typically a logit function) and ε is a normally distributed error term. Then, for multiple stable states, we use a mixture of such regressions:

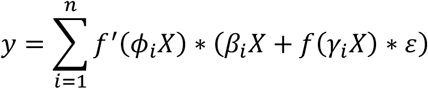

where *ϕ*_i_X is a weighting term with *ϕ* being the vector of parameters, and n is the number of distributions in the mixture. By making the weighting dependent on predictors, we effectively enable stable states to transition into and out of existence along the predictors (consequently the number of stable states can change along a climatic gradient).

The number of distributions in the mixture model does not have to (and typically does not) correspond to the number of stable states in system state variable. This is because stable states of the system state variable are not mean values of underlying distributions but modes of the overall mixture (compound distribution). A mixture of multiple distributions can thus describe one stable state with a wide basin. Therefore, the number of distributions used in the mixture ought to be higher than expected number of stable states of the system. Distributions which do not increase the data likelihood, will have the weighting term close to zero thus will be effectively suppressed. The number of distributions for a particular dataset can be determined using Watanabe-Akaike information criterion (Watanabe, 2013).

The fitted model is then used to infer stable states, computed as local maxima in the compound distribution, and tipping points, computed as local minima of the compound distribution. For specified predictor values, a compound probability density is computed as a weighted sum of the probability density of all distributions in the mixture. We compute the compound probability density for 1,000 equidistant points in the range of the response variable (i.e., the system state variable) for each of 500 equidistant points in the range of a predictor. Consequently, using the compound distribution, we can compute, for each observation, various resilience-related characteristics such as ‘distance to stable state’ or ‘distance to tipping point’ (precariousness).

The combination of probability densities of several distributions can lead to multiple local minima and maxima in the stability landscape. Some of these can be only small bumps which are unlikely to represent real stable states or tipping points, and which have large uncertainty regarding their existence and/or position in the landscape. For calculation of some derived characteristics such as distance to closest stable state or tipping point, it is preferable to avoid these local minima and maxima. To do so, we scale the probability density for each stability curve to be from 0 to 1 and consider as stable states/tipping points only such local minima and maxima which differ in scaled density by at least 0.1. To check the robustness of the results, we also explore thresholds 0, 0.2, and 0.3.

### Dataset

To illustrate the use of the model, we selected one published case of multiple stable states – tree cover in tropical regions, which was described to have three stable states: forest, savanna, and treeless landcover class (Hirota et al., 2011). Whereas the original study explored tree cover along a precipitation gradient on three continents, here we selected only South America since it was found to have the most pronounced stable states. We obtained the original data and used our model to identify stable states along a precipitation gradient. Precipitation and tree cover data were provided by the authors of the original study (Hirota et al., 2011) with primary sources being Mitchell and Jones (2005) and Hansen et al. (2003) respectively.

To show the ability of the model to control for multiple covariates, we also ran the model with added precipitation variability as a second predictor. Precipitation variability is considered an important factor governing the state of ecosystems, especially in water limited systems (Berdugo et al., 2022), which, although not dominant, are present in the tropical zone of South America (Northeast Brazil). For this, we used the BIO15 variable from the WorldClim dataset (Fick & Hijmans, 2017).

### Model specification

To estimate stable states in tree cover along the precipitation gradient, we modelled tree cover as beta distributed. Beta distribution is a continuous distribution ranging from 0 to 1, thus corresponding to proportion of tree cover. Since the beta distribution is not defined for the extreme values 0 and 1 which are common in tree cover data, we transformed the tree cover data to be in range from 0.01 to 0.99 (by multiplying them by 0.98 and adding 0.01). Following the original study, we also excluded observations with mean annual precipitation higher than 3700 mm per year because these were too sparse to reliably estimate stable states. This excluded 66 of 19,371 observations. From the resulting 19,305 observations, we randomly selected 6,000 which we used for the analyses.

Based on Watanabe-Akaike information criterion, we used 7 distributions in the mixture. We modelled their mean, precision and probability as dependent on precipitation. We standardized the predictor to 0 mean and standard deviation 1. We used 0 centred normal prior with precision 0.1 for most parameters. For *ϕ* (parameter of the weighting term), we used a precision of 0.001, and for β_i_ such that i > 2 (parameter of the mean term), we used gamma priors with 0.001 shape and rate. We ran 2 chains with 10,000 iterations and 5,000 burn-in.

### Power analyses

To assess power of the model and to give guidance regarding the required sample size for identification of stable states in a system, we performed power analyses. We generated data by first assigning each observation randomly to one of two states (each with 50% probability), generated predictor values from a uniform distribution ranging from 0 to 1, and system state variable from a normal distribution with the mean depending on the assigned state and standard deviation 1. We changed sample size, distance between the states, and shift – the overlap of those states along the predictor by increasing predictors values for one of the states:

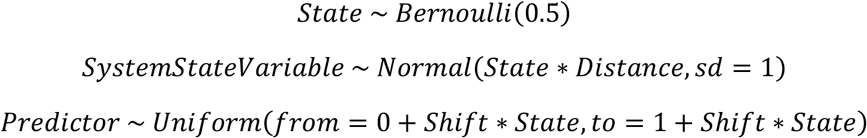

where Distance and Shift are prespecified scalar values. For each combination of parameters, we ran 100 models and assessed power as proportion of models in which an alternative stable state was identified for all observations (in case of Shift > 0 only for observations with overlapping predictor values).

## Results

We identified three stable states in tree cover along a precipitation gradient in the study region (Fig. 2) using mixture model with seven beta regressions (Fig. 3). At low precipitation, there was only treeless state which ceased to exist around 1300 mm/yr. At precipitation 900 mm/yr, started savanna state with tree cover around 20% which existed up to precipitation 2200 mm/yr. At precipitation around 1500 mm/yr, the forest state appeared with tree cover around 80% and was present up to the upper range of the precipitation gradient at 3700 mm/yr (Fig. 2). Therefore, there were two alternative stable states in precipitation ranges approx. 900-1300 mm/yr and 1500-2200 mm/yr. There was a considerable variability around these stable states, especially in relation to the savanna state, but the uncertainty about the position of all three stable states was low (Fig. S1). These stable states were identified irrespectively of the used threshold value (Fig. S2).

**Figure 2:**
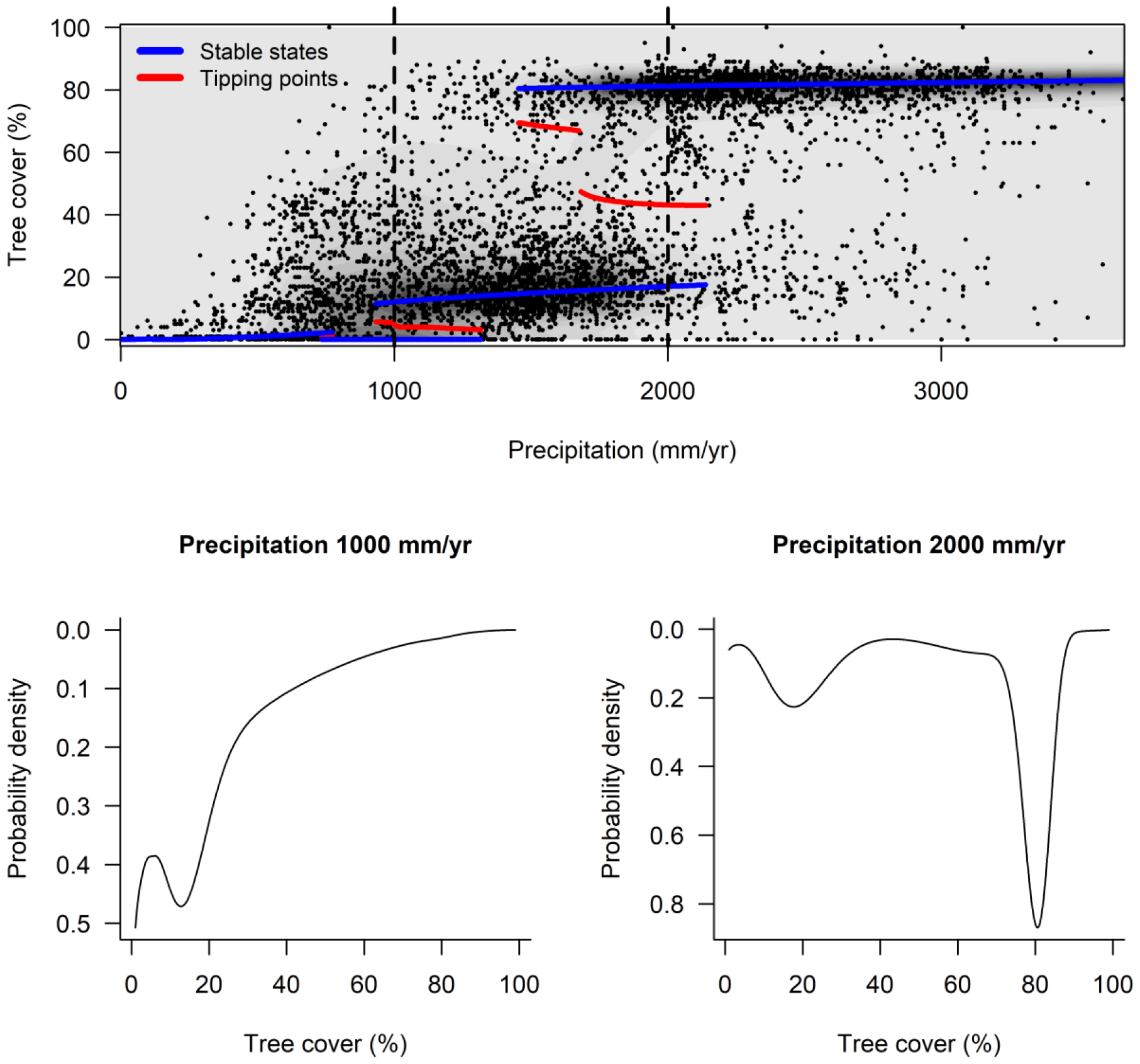
Stable states in tree cover along precipitation gradient (upper part) and stability curves for selected precipitation values (lower part). Upper part: blue lines denote putative stable states, red line denotes tipping points, dashed vertical lines mark selected precipitation values for which stability curves are shown. Shading corresponds to probability density (scaled).

**Figure 3:**
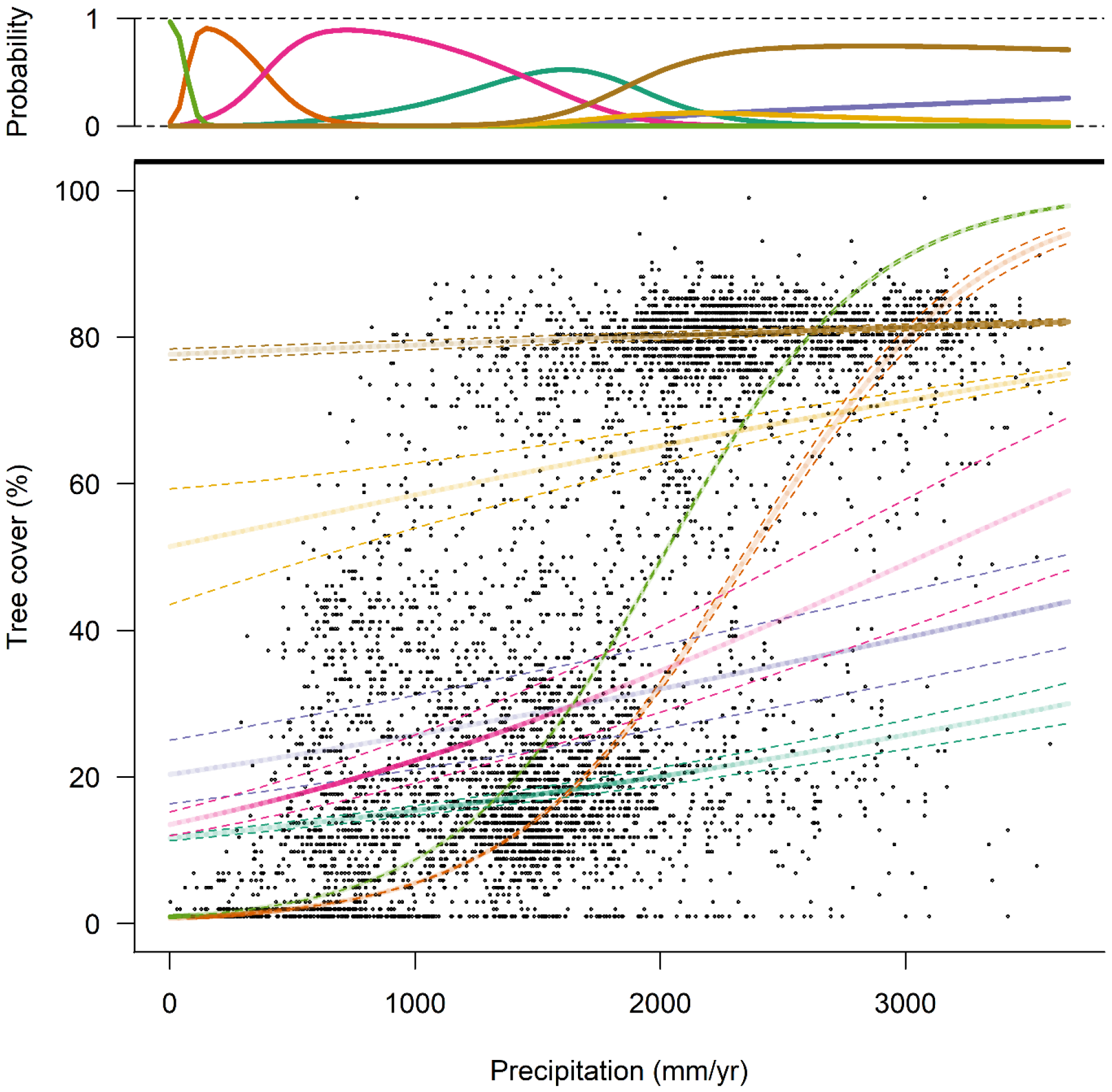
Seven fitted beta distributions along the precipitation gradient. Posterior means are visualized; solid lines denote the mean of each distribution; dotted lines show standard deviation. Upper part of the figure shows probability of each distribution (the weighting term). Opacity of a particular distribution in the lower plot corresponds to the weighting.

The area in the domain of the forest state (i.e., observations in the basin of forest state in stability landscape; 40.3% of the total area) was mostly located in the Amazon basin (Fig. 4). Areas identified as the forest state had mostly high estimated probability density and were near to the stable state in terms of tree cover (Fig. 5). However, 30.3% of the forest state was found to have an alternative stable state (savanna; Fig. 4). The area in the domain of the savanna state (37.4% of the total area) was in southeast and around the Amazon basin. It had a variable estimated probability density and distance to stable state (Fig. 5). There was an alternative stable state for most of this area – for 53.5% it was forest and for 31.5% it was the treeless state (Fig. 4). The area in the domain of the treeless state (22.3% of the total area) was in the southwest of the continent and eastern Brazil. It was mostly close to the stable state, and it had variable estimated probability density (Fig. 5). For 8.5% of this area, there was an alternative stable state savanna. Overall, these patterns were not sensitive to inclusion of precipitation variability as another predictor (Fig. S3).

**Figure 4:**
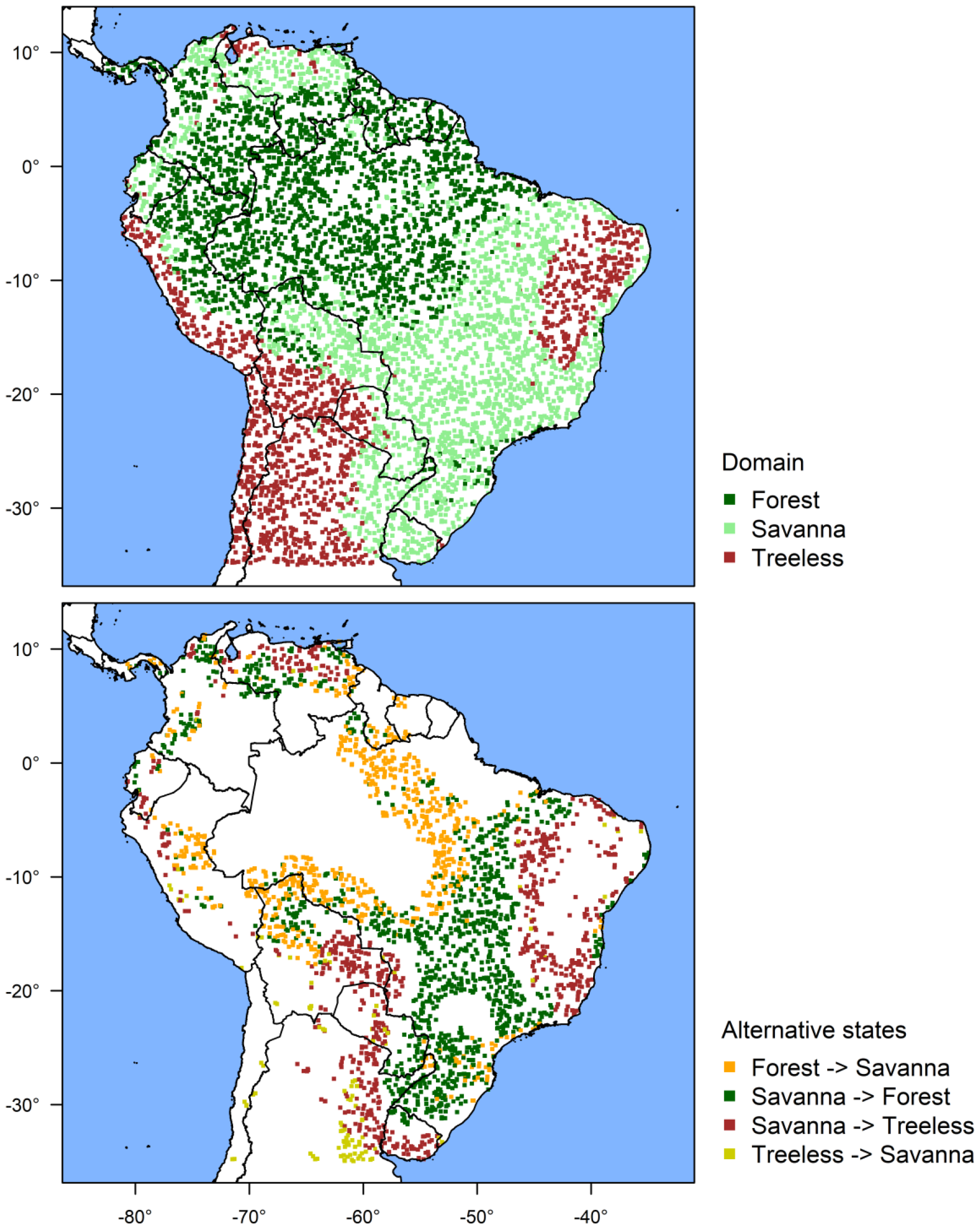
Domains and alternative stable states. Domain is the current stable state for each observation (basin in the stability landscape). Part of the area has an alternative stable state (bottom) given the precipitation values – for forest, it is savanna (30.3% of forest, orange); for savanna it is either forest (53.5% of savanna, green) or treeless state (31.5% of savanna, brown), and for treeless state it is savanna (8.5% of treeless, yellow).

**Figure 5:**
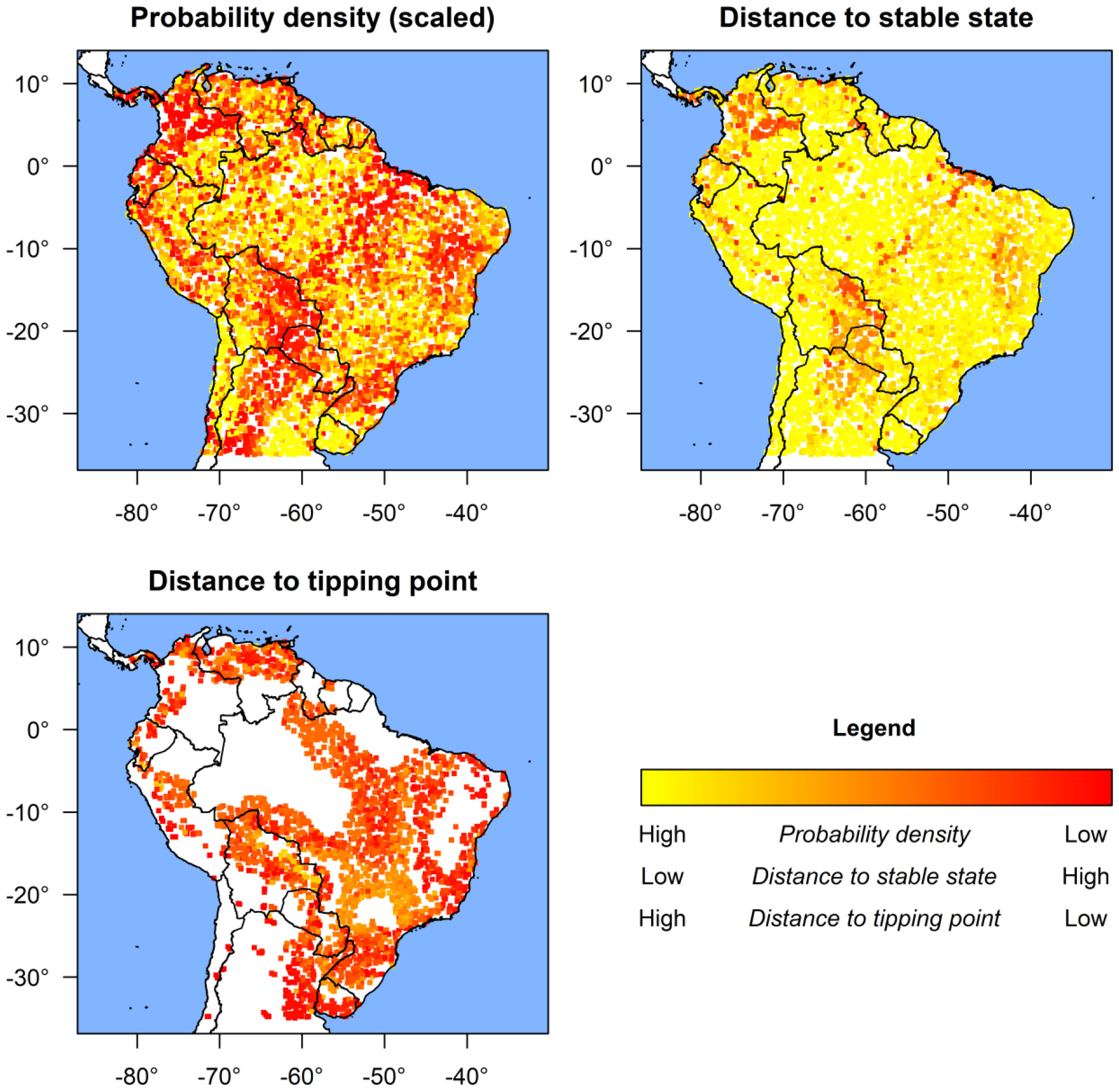
Maps of probability density, distance to the closest stable state and tipping point. There is an alternative stable state and thus distance to a tipping point in 45.9% of the area. Probability density is the estimated probability density in the stability landscape scaled to 0-1. Distance to stable state or tipping point is a difference between current tree cover at the location and tree cover which is estimated to be the closest stable state/tipping point for given precipitation.

Power analysis showed that a sample size of at least 200 was necessary using this model for reliable identification of alternative stable states. In order to be identifiable as distinct, distances between stable states had to be at least 4 standard deviations from each other, and the stable states could overlap only little along the predictor gradient at the expense of slight decrease in the power (Fig. 6).

**Figure 6:**
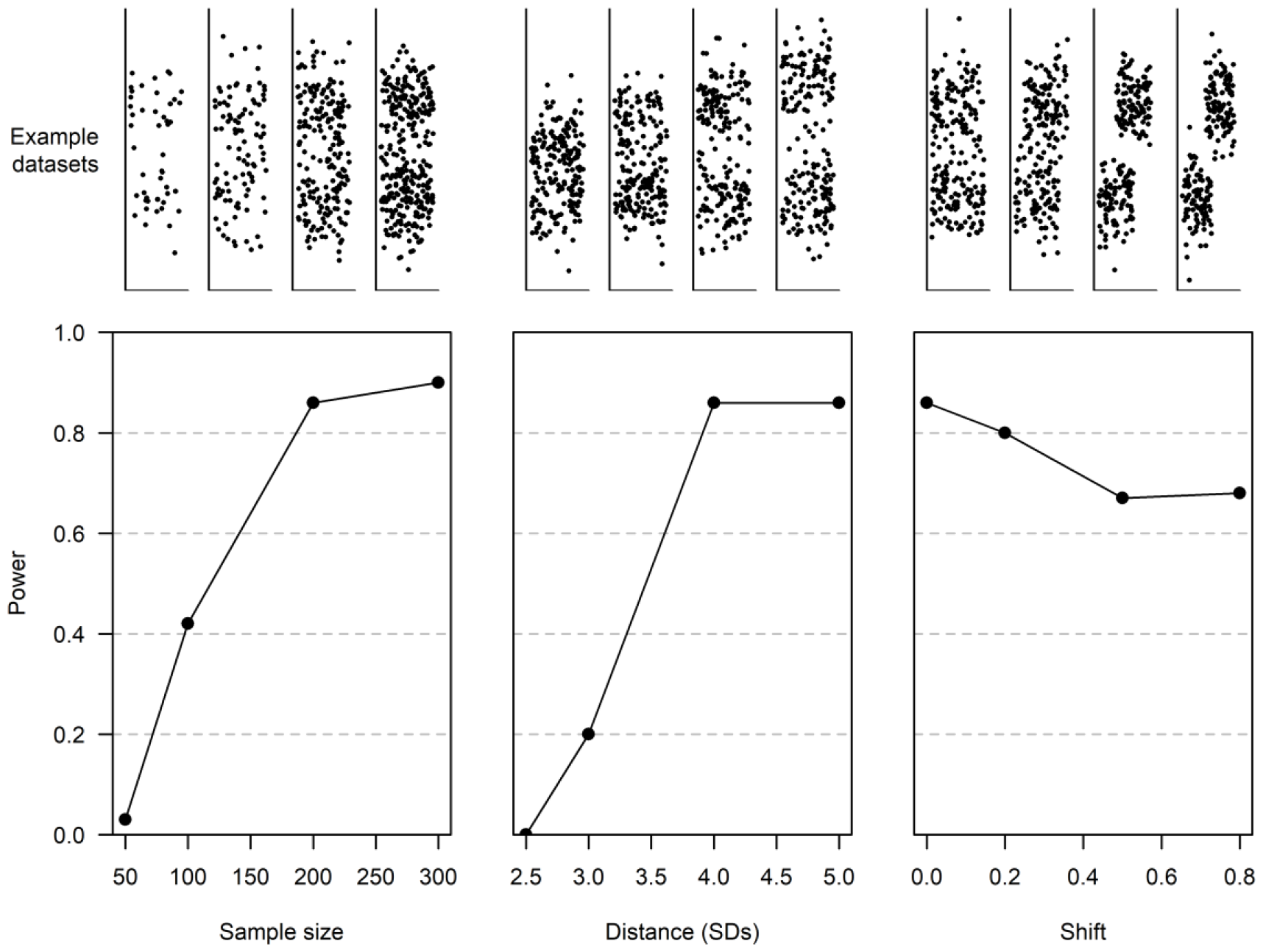
Power analysis for different sample sizes, distances of stable states, and shift between stable states along the predictor. In analyses of different sample sizes, distance was 4 standard deviations and there was no shift; in analyses of different distances, sample size was 200 and there was no shift; in analyses of different shifts, sample size was 200 and distance was 4 standard deviations. Each example dataset corresponds to the point below it.

## Discussion

We used our mixture model to identify previously described stable states in tropical tree cover in the neo-tropics and assess the resilience of these locations. The original study (Hirota et al., 2011) described three stable states – forest, savanna, and the treeless state, centred around tree cover values of 80%, 20%, and 0%, respectively. Our model successfully captured these states. Moreover, we show which areas have alternative stable states and map various resilience metrics (Fig. 4, 5) suggesting areas prone to change. Specifically, the savanna state has an alternative stable state in form of the treeless state or forest along most of its precipitation range representing 85% of its area. A large part of this area has low estimated probability density and is close to a tipping point highlighting potential for regime shifts. To a lesser degree, the same is true for the forest state. Although savanna is present as an alternative state to it only for minority of its precipitation range, this represents nearly one third (30.3%) of total forest area. Under current precipitation levels, the treeless state does not seem to be prone to change into the savanna (apart from rather small area at the southern limit of studied range).

There are three main points in which our model improves the original approach to identify stable states from a distribution of ecosystem state variable. Firstly, the original approach has an important assumption which we relaxed. It assumes that system stable states have a constant value of the system state variable along a climatic gradient and thus stable states are identifiable based on overall distribution of ecosystem state values (Hirota et al., 2011; Scheffer et al., 2012). However, the stability of a system state refers to an equilibrium among the components of the ecosystem. It does not require that value to be constant across different external conditions (e.g., along a climatic gradient). As stable states are often kept in place by the interaction of multiple drivers (such as herbivory intensity, fire frequency, population dynamics; van Langevelde et al., 2003) and these drivers are likely climate dependent (Staal et al., 2018), we can expect a system stable state value to change along a climatic gradient. Therefore, the identification of stable states should take the climatic gradient(s) into consideration, e.g., as predictor(s) in a mixture regression model.

A second important difference between our model and the original approach is that the latter used kernel smoothing when exploring stable states along the precipitation gradient, which requires an arbitrary choice of the smoothing window size. In contrast, our approach could be used to model the generative process behind the observed distribution of the system state variable. Identified distributions could represent effects of components of the system – ecological drivers or their combination which “push” the system to a particular value of system state variable. For example, in a growth-herbivory model (Fig. S4), growth would push the system towards carrying capacity and herbivory towards zero. Depending on the relative strength of these two components, the system can have zero, one, or two stable states between the extreme values (Fig. S4). In the mixglm model, the mean of a distribution corresponds to the value towards which the ecological driver “pushes” the system, the probability to its power, and the variation to decay of its influence with distance. However, caution is necessary when interpreting the distributions – estimation of mixtures can be challenging with different combinations of distributions approximating the same stability landscape. Therefore, a strong a priori knowledge of drivers of the system is required to interpret the distributions as effects of ecological drivers. With such knowledge, the model could be used to test hypothesized drivers of the system, possibly by specifying number of distributions or constraining their parameters based on the hypothesis.

Finally, unlike the original approach, our model follows from well-established regression modelling which allows flexible extensions of the model. The first of such extensions is the option to include multiple predictors, which we illustrate by including precipitation variability into the model of tree cover in South America (Fig. S3). This is important since climate cannot often be characterized by a single gradient in a satisfiable way (Bojinski et al., 2014). There is also a potential to include other terms into the model such as spatial or temporal one to account for spatial/temporal autocorrelation. Yet, fitting of mixture models is non-trivial (Hurn et al., 2003) and including correlation structure into them is a topic for further research.

Using power analyses, we showed that sample size around 200 is necessary to reliably identify alternative stable states even in ideal conditions where they are only two and are well separated. This sample size is easily achievable using remote sensing products but could be more challenging in other datasets where alternative states have been proposed but which are more challenging to generate large datasets (e.g., shallow lake productivity; Davidson et al., 2023).

Another requirement is a suitable ecosystem state variable. Since the model identifies stable states for the distribution of this variable, there must be no discontinuities in the distribution caused by its calculation. Therefore, variables (e.g, gross primary production based on MOD17 algorithm; Running & Zhao, 2015) which are calculated using categorical inputs such land cover types might not be suitable. Even without such inputs, complex algorithms can lead to discontinuities or decreased resolution in part of the distribution of ecosystem state variable which might limit the interpretation. This is also the case for tree cover (Hansen et al., 2003) which has been shown to not be suitable for small geographical scales and have worse resolution in low tree cover values (Hanan et al., 2014, 2015; Staver & Hansen, 2015).

Apart from the model itself, we have implemented a series of functions to process the model’s results. Model parameters are used to compute a compound distribution which, when evaluated for different precipitation values, create the stability landscape (Fig. 2). The stability landscape can consequently be used to extract various resilience characteristics (e.g., probability density meaning how likely is the observation to change its system state value, distance to stable state capturing difference between the system state value of the observation and its estimated stable state, and distance to tipping point being the necessary change in system state value to transition into a domain of different stable state) for each observation and map them (Fig. 4), which strongly enhances interpretability of results. Finally, the model can be easily used for predictions of future stability landscapes, resilience metrics, and stable states (Fig. 7).

**Figure 7:**
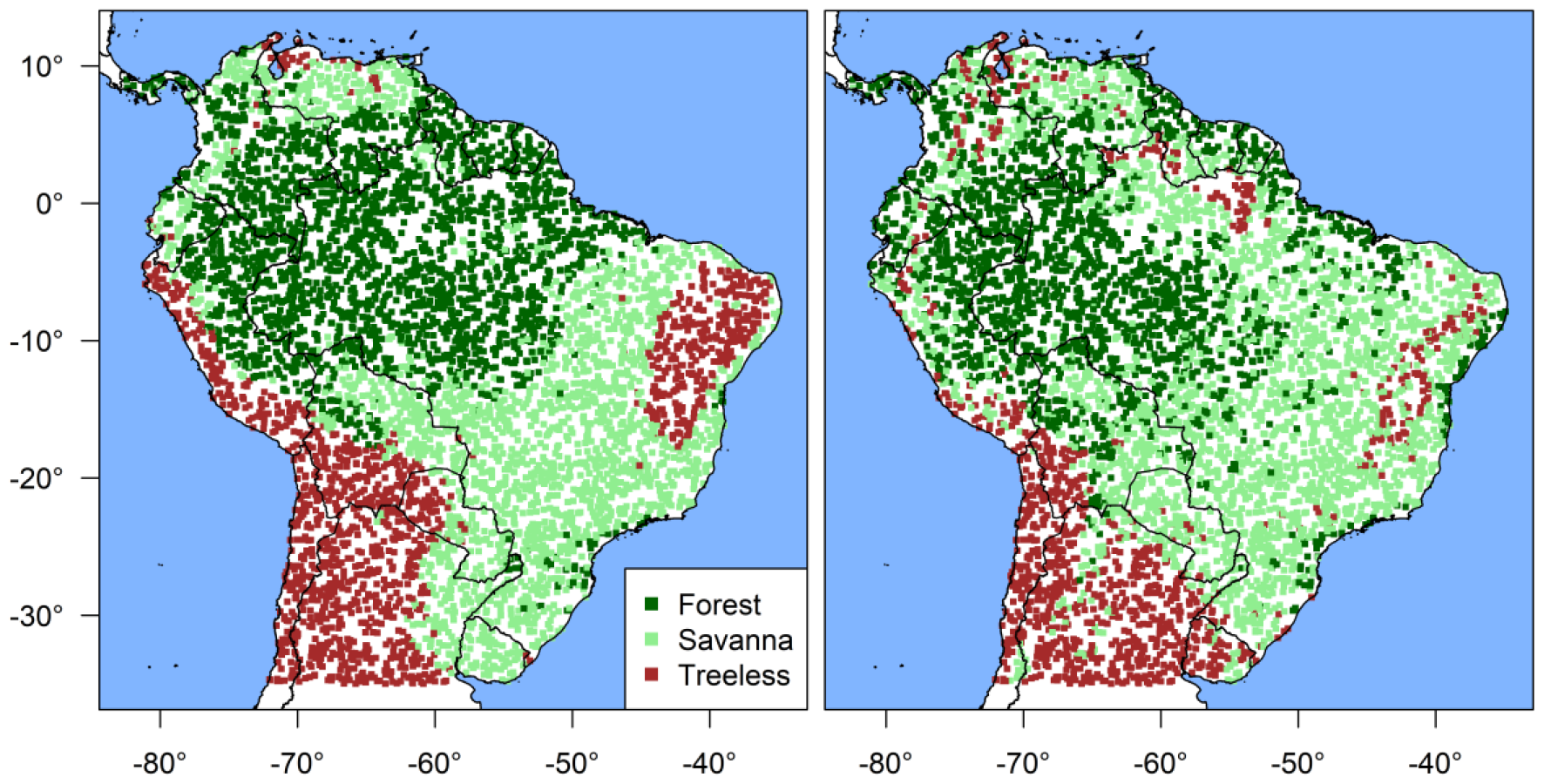
Estimated (left) and predicted (right) domains of stable states. Prediction is based on precipitation from CORDEX South America model, scenario RCP 4.5 (intermediate scenario) for period 2081-2100 (Gutiérrez et al., 2021; Iturbide et al., 2021). Domain changes are predicted for 30.0% of area with 15.1% of area shifting towards less forested states.

## Conclusion

We present an implementation of a model to estimate system stable states and their resilience from the distribution of a selected system state variable (e.g., tree cover), employing one or more explanatory covariates such as climatic variables (here, annual precipitation and precipitation variability). The model was able to identify previously published stable states in tree cover of the tropical region of South America. Furthermore, it is more flexible than its predecessor in what phenomena it can describe, it is less strict in its assumptions, and it is extendable to account for more complex data structures. As such, it promises possibility for further development and applications. The model and associated tools are available as an R package “mixglm” at https://github.com/adamklimes/mixglm.

## Acknowledgements

We are grateful to Marina Hirota for providing data on tree cover and precipitation of tropical regions. The work received funding from the Research Council of Norway (grant #320602).

## Authors’ contributions

JDC, AWRS, and MMF came up with the idea; all authors conceived the research; JDC and AK prepared the model and analyses; AK wrote the manuscript with contributions of all other authors.

