## Supplementary information for "[mixglm]: A tool for modelling ecosystem resilience"

Content:

- Figure S1-4

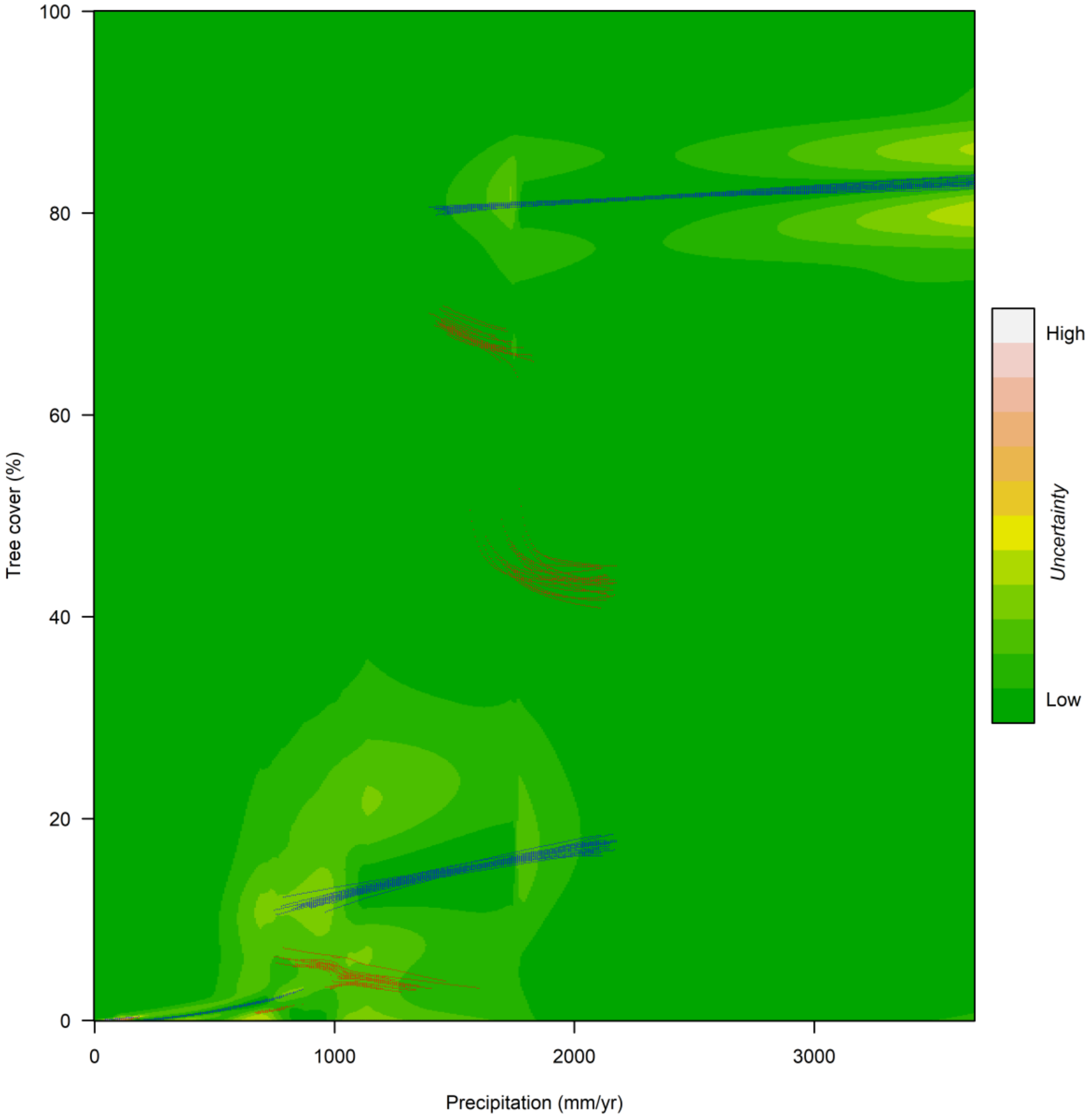


*Figure S1: Uncertainty of the stability landscape. Standard deviation of scaled probability density of observations based on 20 random posterior samples is visualized along estimated stable states (blue lines) and tipping points (red lines) for each of these samples. Lighter colours denote higher uncertainty of the probability density. Position of stable states and tipping points is very similar across all plotted samples.*

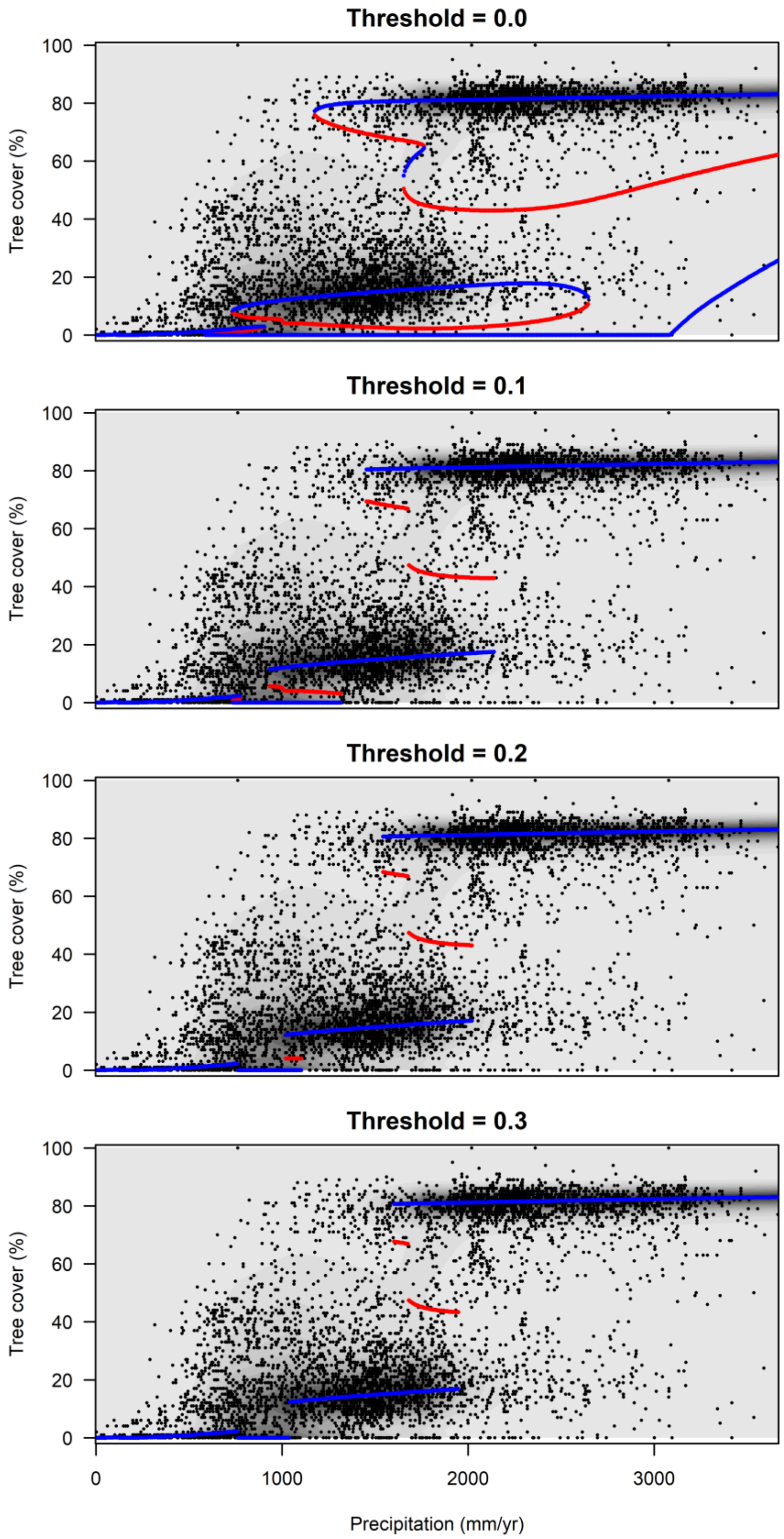


*Figure S2: Different thresholds for stable states and tipping points in tree cover along precipitation gradient. Blue lines denote putative stable states, red line denotes tipping points. Shading corresponds to probability density (scaled). For local minima and maxima of probability density to be considered stable states and tipping points respectively, their scaled probability density has to differ at least by the threshold value. The three states – forest, savanna, and treeless state – are consistently identified as stable states irrespective of the threshold value.*

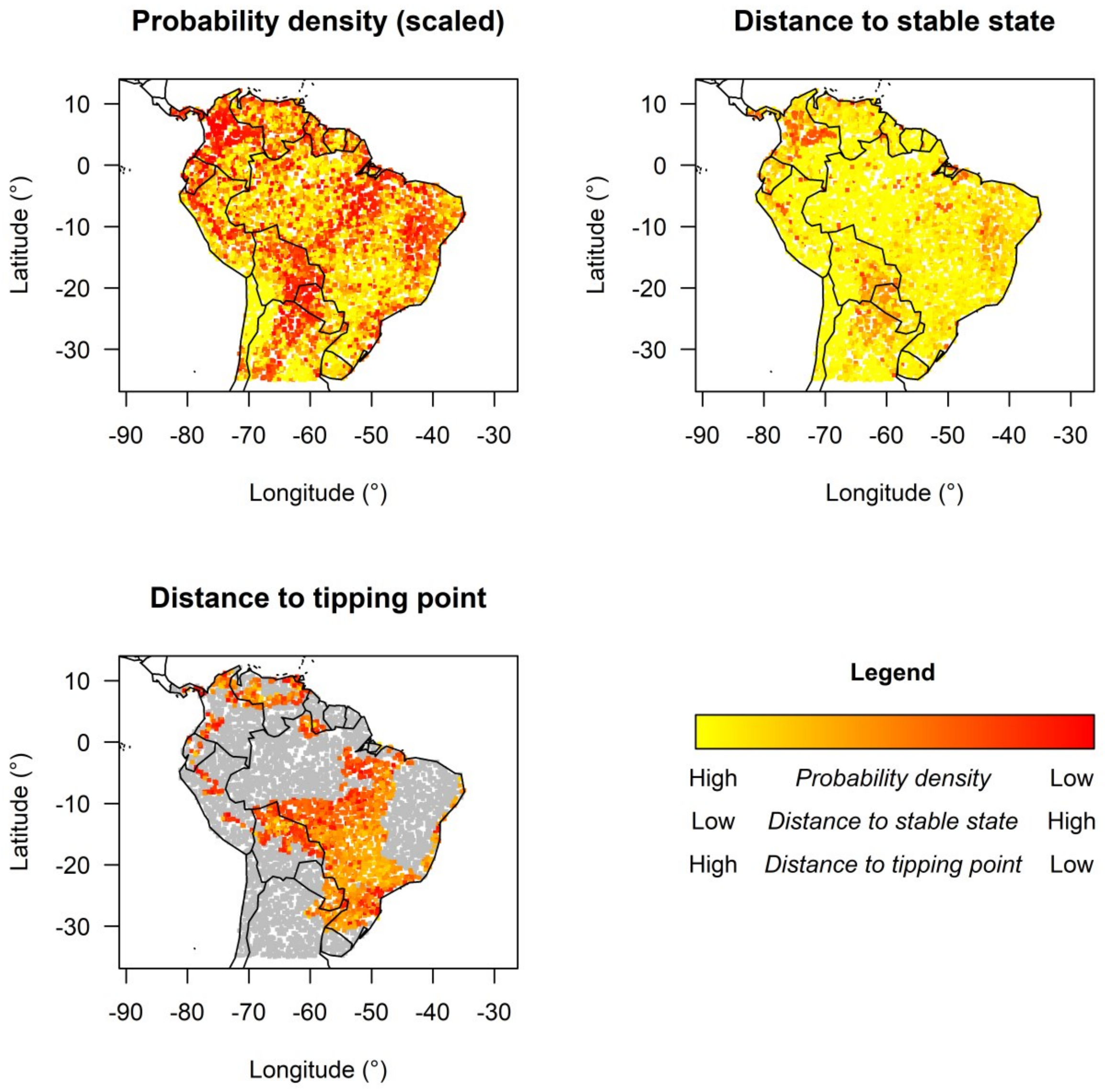


*Figure S3: Maps of probability density, distance to the closest stable state and tipping point for the model of tree cover including both precipitation and precipitation variability. Grey colours denote no tipping point. Probability density is the estimated probability density in the stability landscape scaled to 0-1. Distance to stable state or tipping point is a difference between current tree cover at the location and tree cover which is estimated to be the closest stable state/tipping point for given precipitation. Patterns are very similar to results of the model without precipitation variability (Fig. 5).*

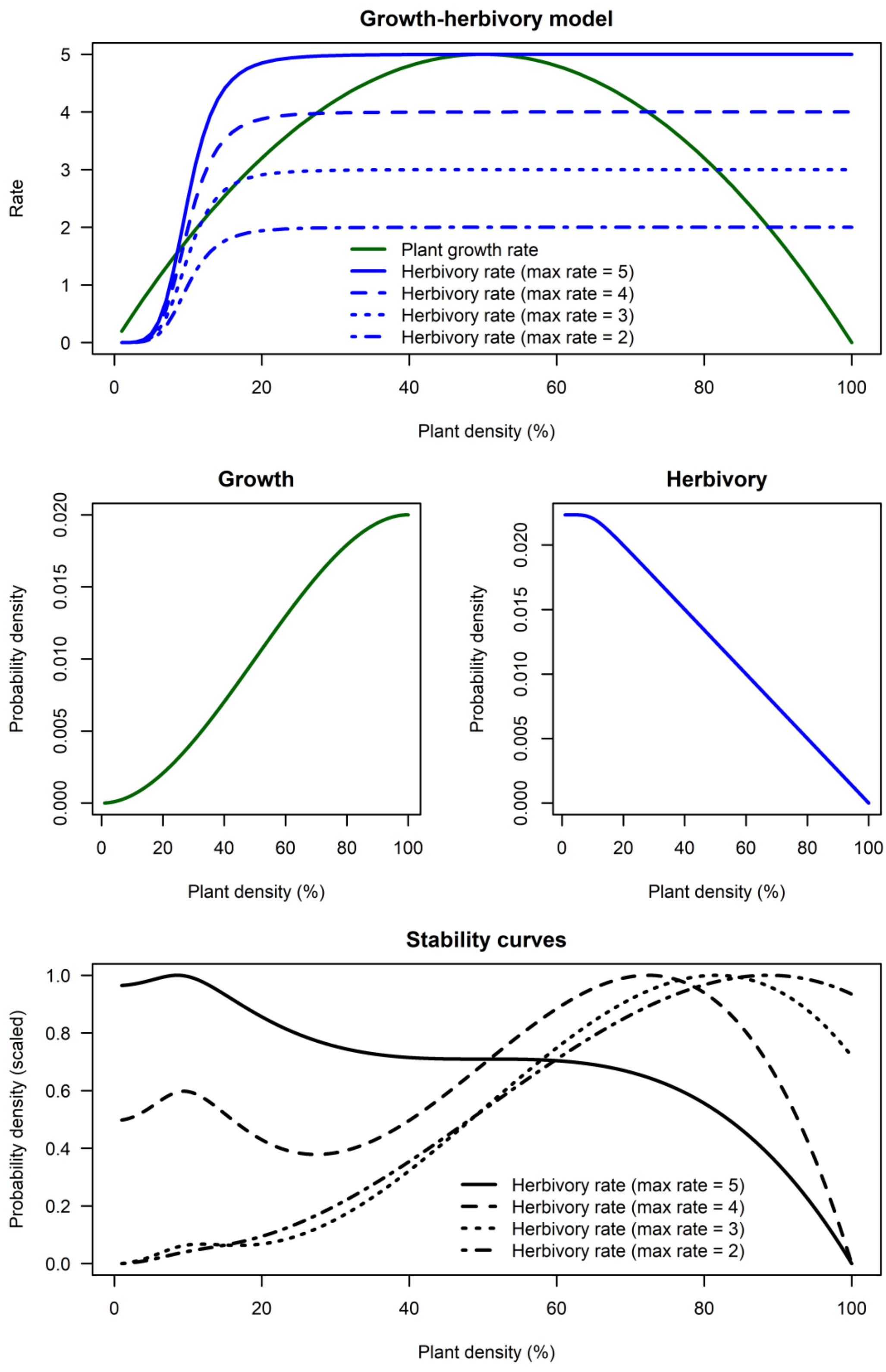


*Figure S4: Simple growth-herbivory model illustrating that combination of two components of the system can lead to stability curve with one or two stable states. Top: Plant growth rate and herbivory rate can intersect at one or more points along plant density gradient depending on maximum herbivory rate. Middle: Slope of the distribution is proportional to growth/herbivory rate (the same shape for all herbivory rates once standardized). Bottom: Combined effects of both components create stability curves with minima and maxima corresponding to intersections of growth and herbivory rates as seen in the top part of the figure.*
